# Divergence in poxvirus-encoded E3-like proteins can dictate poxvirus activation of cellular necroptosis

**DOI:** 10.1101/2024.12.05.627069

**Authors:** Junior A. Enow, Saige Munig, Mathew L. Sample, Gil Speyer, Raghavi Chandrakanth, Jacqueline Williams, James Bonner, Grant McFadden, Bertram Jacobs, Masmudur M. Rahman

**Affiliations:** Molecular and Cellular Biology Program, Arizona State University; School of Life Sciences, Arizona State University; Center for Personalized Diagnostics, Biodesign Institute; Biodesign Center for Molecular Design and Biomimetics Biodesign Institute; ASU-Banner Neurodegenerative Disease Research Center Biodesign Institute; Research Technology Office, Arizona State University, Tempe, Arizona 85287

## Abstract

Poxviruses encode a plethora of proteins to modulate diverse cellular responses against viruses. Poxvirus-encoded E3-like proteins are multifunctional, regulating diverse cellular antiviral responses. The canonical Vaccinia E3-like proteins have two domains: an N-terminal Z-form nucleic acid binding domain (Zα-BD) and a C-terminal double-stranded RNA binding domain (dsRNA-BD)-.Using protein sequence and structural homology modeling, we identified the presence of dsRNA- BD-containing proteins in all the poxviruses except Avipoxviruses, Salmon poxvirus and Entemopoxviruses. However, the acquisition of these proteins likely happened under three distinct events. Using structural homology modeling and FATCAT score, we can classify E3-like proteins in three distinct categories: i) the E3-like proteins with highly conserved dsRNA-BD but with or without the N-terminal domain, present in most poxviruses; ii) unconventional E3-like proteins with highly diverged dsRNA-BD, present in Macropoxvirus and Molluscipoxvirus and iii) E3-like protein with dsRNA-BD that may have different origin present in Crocodilepoxvirus.^12–52–6^

Members of Leporipoxvirus, Waddenpoxvirus, Cetaceanpoxvirus, and selected members of Orthopoxvirus contain E3-like proteins missing the N-terminal Zα-BD required for necroptosis inhibition. Additionally, using Alphafold, we show that the Zα-BD of Chordopoxviruses E3-like proteins is structurally more variable than the ds-RNA binding domain. Compared to members of Orthopoxviruses–Vaccinia virus (VACV) and Cowpox virus (CPXV) that have been shown to inhibit necroptosis and contain an N-terminus Zα-BD of the canonical E3 protein, our results show that members of leporipoxviruses induce necroptosis in human and mouse necroptosis competent cell lines. Furthermore, myxoma virus (MYXV) infection activates RIP1 and RIP3-mediated necroptosis in both human and mouse necroptosis-competent cells. These data suggest that Leporipoxviruses lack countermeasures to necroptosis compared to Orthopoxviruses that encode multiple key regulators of necroptosis, possibly due to a lack of selective pressure within the viral host species (Lagomorphs).

**Graphical Abstract:** 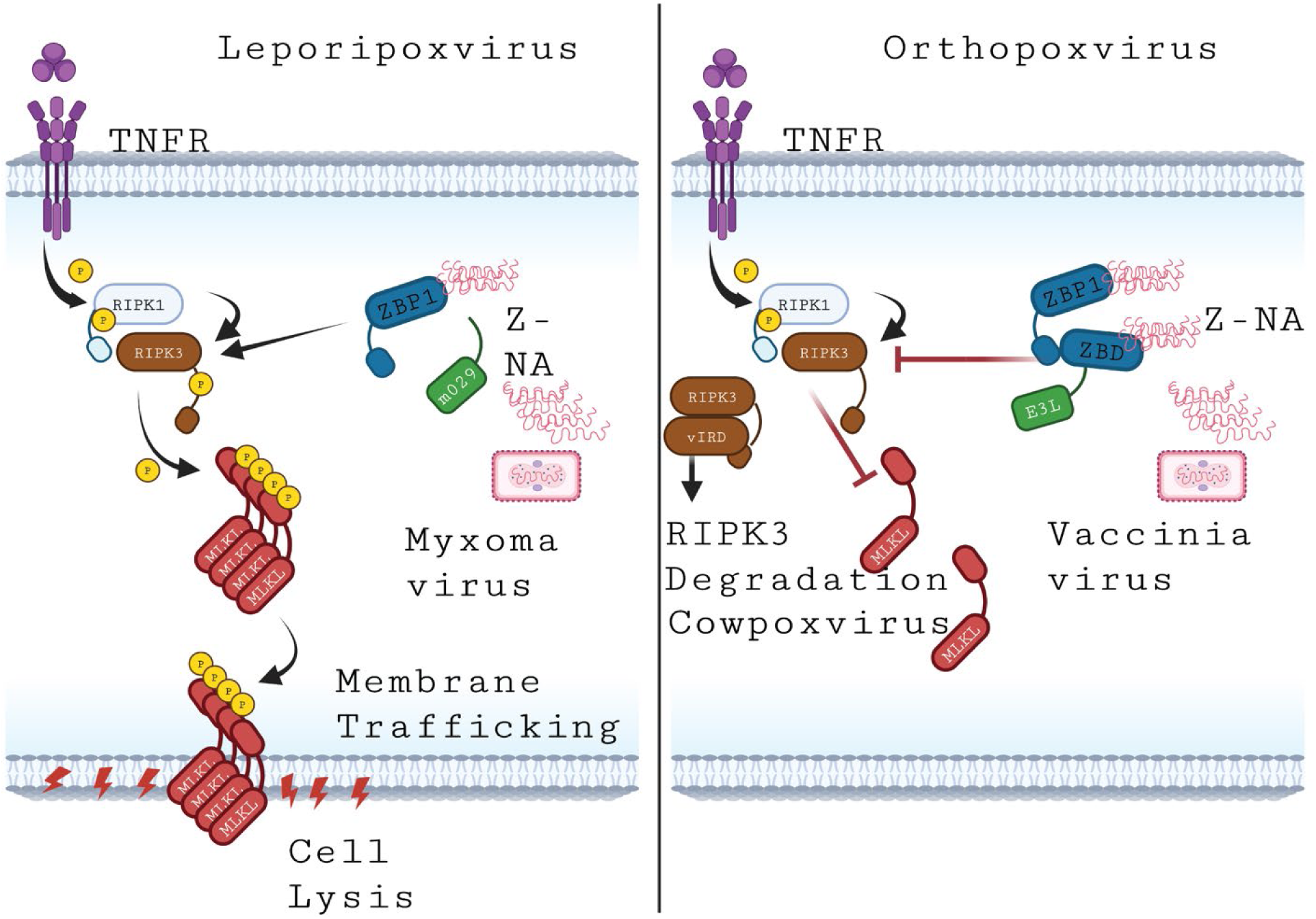

## Introduction

Poxviruses are unique amongst other viruses in that they replicate exclusively in the cytoplasm of infected cells^1,2^. In order to successfully replicate in the cytoplasm of infected cells, poxviruses encode a plethora of immune evasion proteins. Poxvirus-encoded vaccinia virus (VACV) E3-like family of proteins are one of the key immune evasion proteins. E3 has two conserved domains: the N-terminal Z-form nucleic acid binding domain (Zα-BD) and a C-terminal double-stranded RNA binding domain (dsRNA-BD). The C-terminal dsRNA-BD is required for the sequestration of dsRNA to avoid the activation of dsRNA-dependent antiviral enzymes such as protein kinase R (PKR)^3–7^. The N-terminal domain on the other hand is required for the inhibition of virus-induced necroptosis^4,8^. The dsRNA-BD present in the E3 family of proteins displays a higher level of sequence similarity than the N-terminal domain. Studies from different poxviruses suggest that dsRNA-BD is essential for the in vitro replication and in vivo pathogenesis of poxviruses^4,8–10^. This conservation in dsRNA-BD was further confirmed by observations that a dsRNA-BD containing protein from a completely different virus such as influenza virus encoded NS1 protein can rescue *E3L*-lacking VACV replication in cell culture^11^. However, the N-terminal domain is dispensable for replication in most cell culture settings but is required for pathogenesis in mice^4,6,8,10^. Furthermore, the N-terminal domain is partially or completely missing in some members of poxviruses^5,7^. This N-terminal diversity among the E3 family of proteins suggests that the E3 family of proteins might have adapted host-specific immune regulatory functions for in vivo pathogenesis in selected hosts. It is also possible that the E3 protein family are involved in regulating additional cellular pathways that are yet to be discovered.

Necroptosis is a caspase-independent programmed cell death characterized by cell swelling, membrane rupture, and the release of pro-inflammatory molecules^12^. Receptor Interacting Protein Kinase-3 (RIPK3) and mixed-lineage kinase domain-like protein (MLKL) are the main drivers of necroptosis^12^. Upon phosphorylation of RIPK3 by upstream stimuli, RIPK3 phosphorylates MLKL, and MLKL oligomerizes to the plasma membrane and terminates in cell lysis^13^. Necroptosis can be activated via the extrinsic or the intrinsic pathway^14^. Extrinsic necroptosis is mediated by death receptors and their ligands like TNF-alpha and FasL ligand-receptor interactions; this cascade leads to the phosphorylation of RIPK1 and RIPK3 and the later phosphorylation of MLKL^12–14^.

Phosphorylated MLKL oligomerizes and localizes to the plasma membrane, leading to rupture^14^. Since necroptosis-mediated activation of cell death is a cellular defense mechanism against viruses, some virus-encoded proteins can regulate necroptosis^4,8^. For example, the N-terminal domain of VACV E3 protein^4,8^. Poxviruses also encode additional proteins, such as vIRD (viral inducer of RIPK3 degradation) and vMLKL (viral MLKL) to regulate cellular necroptosis^15,16^.

We previously showed that specific mammalian lineages, including Leporids, cetaceans, and some rodents, contain inactivating mutations in either RIPK3 or MLKL that render necroptosis non-functional in these species^3^. Simultaneously, we demonstrated a correlation between the lack of N-terminal domain of E3-like proteins required for necroptosis inhibition amongst Leporipoxviruses^3^. However, members of leporipoxviruses were not tested for their ability to activate necroptosis in necroptosis-competent cells.

Here, we identify two unique dsRNA-containing proteins within Chordopoxviruses, in addition to the traditional E3 protein. We show that the Zα-BDs of Chordopoxviruses E3 proteins are structurally more variable than the ds-RNA binding domain. The alpha-fold structure of E3 proteins from Poxviruses lacking the Zα-BDs show unique modifications at the N-terminus region.

Compared to VACV and CPXV, which have been shown to inhibit necroptosis^4,15^. Members of leporipoxviruses activate necroptosis in necroptosis-competent human and mouse cells.

Furthermore, MYXV (a model Leporipoxvirus) infection activates RIPK1 and RIPK3-mediated necroptosis in these cells, suggesting that Leporipoxviruses lack a countermeasure to the activation of cellular necroptosis.

## Results

### E3-like proteins encoded by the members of Chordopoxviruses can be classified into three families based on the origin of structurally conserved C- terminus dsRNA-BD

Poxvirus E3-like proteins are indispensable for virus replication and pathogenesis amongst poxviruses^6,17,18^. To understand the conservation of the E3-like proteins across poxviruses, we performed a sequence homology search using the Basic Local Alignment Search Tool (BLAST) protein-protein BLAST and (Domain Enhanced Lookup Time Accelerated) DELTA-BLAST using Vaccinia virus E3 protein as a query against the Poxvirus proteome on the NCBI (National Center for Biotechnology Information) database. Our screen revealed that Avipoxviruses, Salmonpoxvirus, and Entomopoxviruses did not contain an E3-like protein (Figure 1A). Suggesting a different viral protein might be involved in viral nucleic acid regulation in these viruses or the mechanism of dsRNA regulation differs from the canonical model. We modeled the structure of all the E3-like proteins from our initial screen using AlphaFold2. We observed a unique dsRNA-BD-containing protein structure belonging to Macropoxvirus and Molluscipoxvirus, which differs from the canonical E3-like proteins belonging to most Poxviruses (Figures 1B, 1C and 1E). Because of the lack of sequence and structural similarity across all the E3-like proteins in our screen, we asked if poxviruses share a common E3 ancestor.

**Figure 1.**
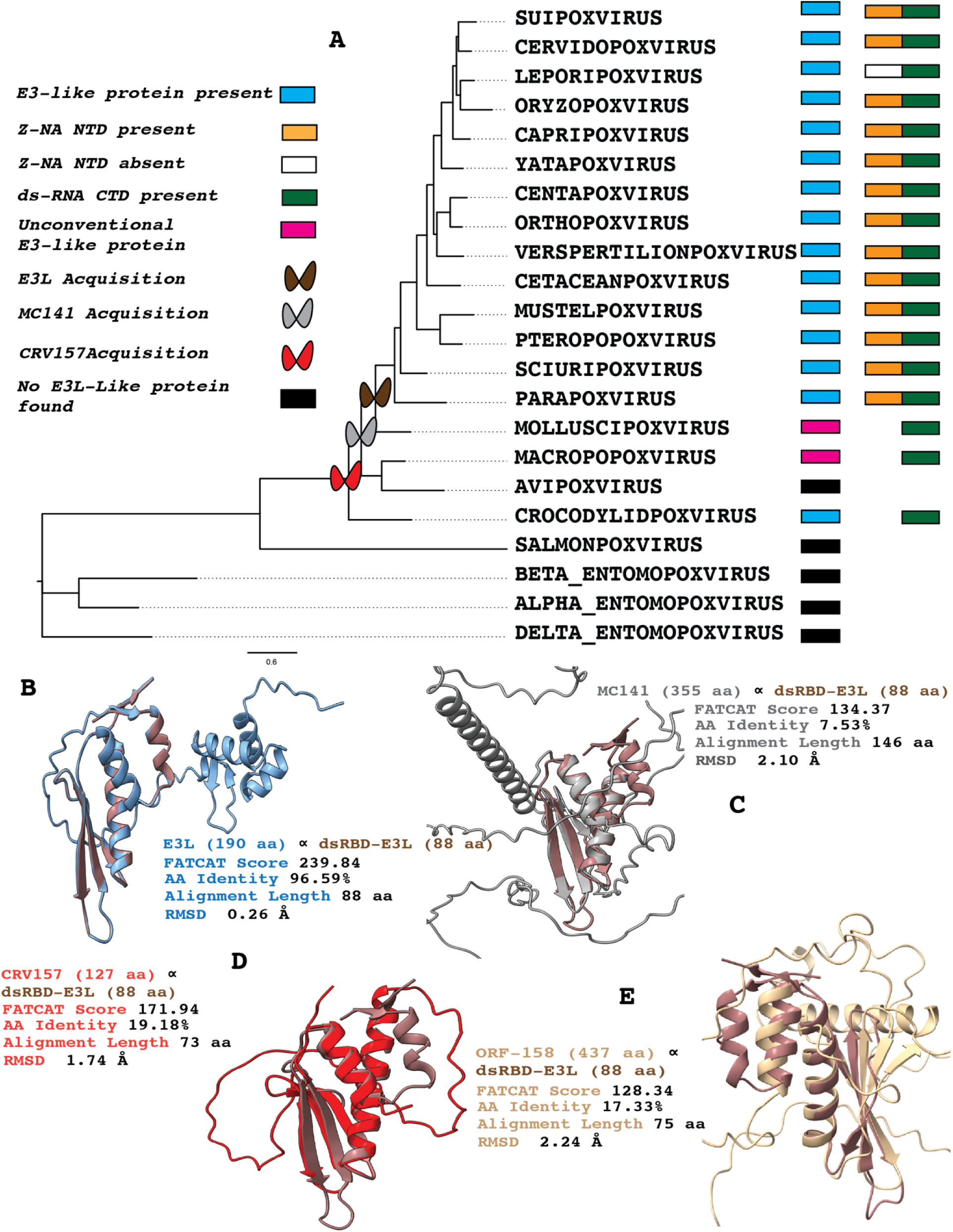
Diversity of double-stranded RNA binding domain proteins within Poxviruses. (A) Phylogenetic tree of all poxvirus families showing the acquisition of dsRNA-BD proteins. Structural homology showing FATCAT score, amino acid identity, alignment length and RMSD value comparing the dsRNA-BD of VACV to (B) VACV E3, (C) Molluscipoxvirus MC141, (D) Crocodilepoxvirus CRV157 and (E) Macropoxvirus ORF-158.

Interestingly, our synteny analysis revealed that Poxvirus dsRNA-BD-containing proteins had 3 distinct acquisition events (Figure 1A and S1A). All Chordopoxviruses except Molluscipoxvirus, Macropoxvirus, Avipoxvirus, Crocodylidpoxvirus, and Salmonpoxvirus share the same ancestral E3- like protein as Vaccinia virus (Orthopoxviruses) (Figure 1A). Molluscipoxvirus and Macropoxvirus share a unique dsRNA-BD-containing protein (MC141 and ORF158) present in no other poxvirus family (Figures 1A, 1C, and 1E). Crocodilepoxvirus contains a dsRNA-BD-containing protein (CRV157) like E3 but with a different origin (Figure 1A, 1D, S1A, and S1B). We used FATCAT structural comparison using the dsRNA-BD of Vaccinia virus E3 as our standard model and compared it with the dsRNA-BD-containing proteins from our screen. Molluscipoxvirus MC141 had a significant FATCAT score of 134.37, an RMSD of 2.10 Å, and an amino acid (aa) identity of 7.53%, suggesting a shared dsRNA-BD homology with VACV dsRNA-BD (Figure 1C).

Crocodilepoxvirus had a significant FATCAT score of 171.94 and an aa identity of 19.18%, an RMSD 1.74 Å (Figure 1D). Macropoxvirus ORF-158 was significantly similar to VACV-dsRNA-BD (FATCAT Score 128.34, 17% aa identity and 2.24 Å RMSD) (Figure 1E). Western Grey Kangaroo Poxvirus gp143 and Saltwater Crocodilepoxvirus (CRV157) were also significantly related to VACV- dsRNA-BD (Figure S1C and S1B).

### Recurrent loss of the Z-form nucleic acid binding domain (Zα-BD) of the E3-like proteins required for intracellular Necroptosis inhibition

Our previous work showed a correlation between the loss of RIPK3 and MLKL among certain mammalian species and the corresponding lack of a Zα-BD of the accompanying infecting poxvirus^3^. We performed phylogenetic analysis using the poxvirus E3-like proteins from our initial screen to understand how the loss of the Zα-BD clusters across different families of Poxviruses.

Phylogenetic tree construction of E3 proteins showed a pattern of loss of the Zα-BD at different Poxvirus families, suggesting that loss at the E3 N-terminus regions are independent events (Figure 2A). Interestingly, we observed that the N-terminus Zα-BD structures of most of the E3 proteins in our screen had additional secondary structures not present within the N-terminus of VACV-E3 (the standard Z-NA model). To understand the extent of the structural diversity occurring at the N-terminus domain of the Poxvirus E3 protein, we performed a structural homology comparison using FATCAT and TM-align. The E3 proteins were broken into the N-terminus Zα-BD and C-terminus dsRNA-BD. Vaccinia virus N and C terminus were used as the query and compared with all Poxviruses that contained an N and C terminus Zα-BD and dsRNA-BD, respectively (Figures 2B and 2C). Our results show a lower FATCAT score and higher RMSD value for the N-terminus domain than the C-terminus, indicative of lower structural similarity at the Zα-BD domain (Figures 2D and 2E). The lower structural homology at the N-terminus domain of E3 suggests modifications to the Zα-BD that might alter its function.

**Figure 2.**
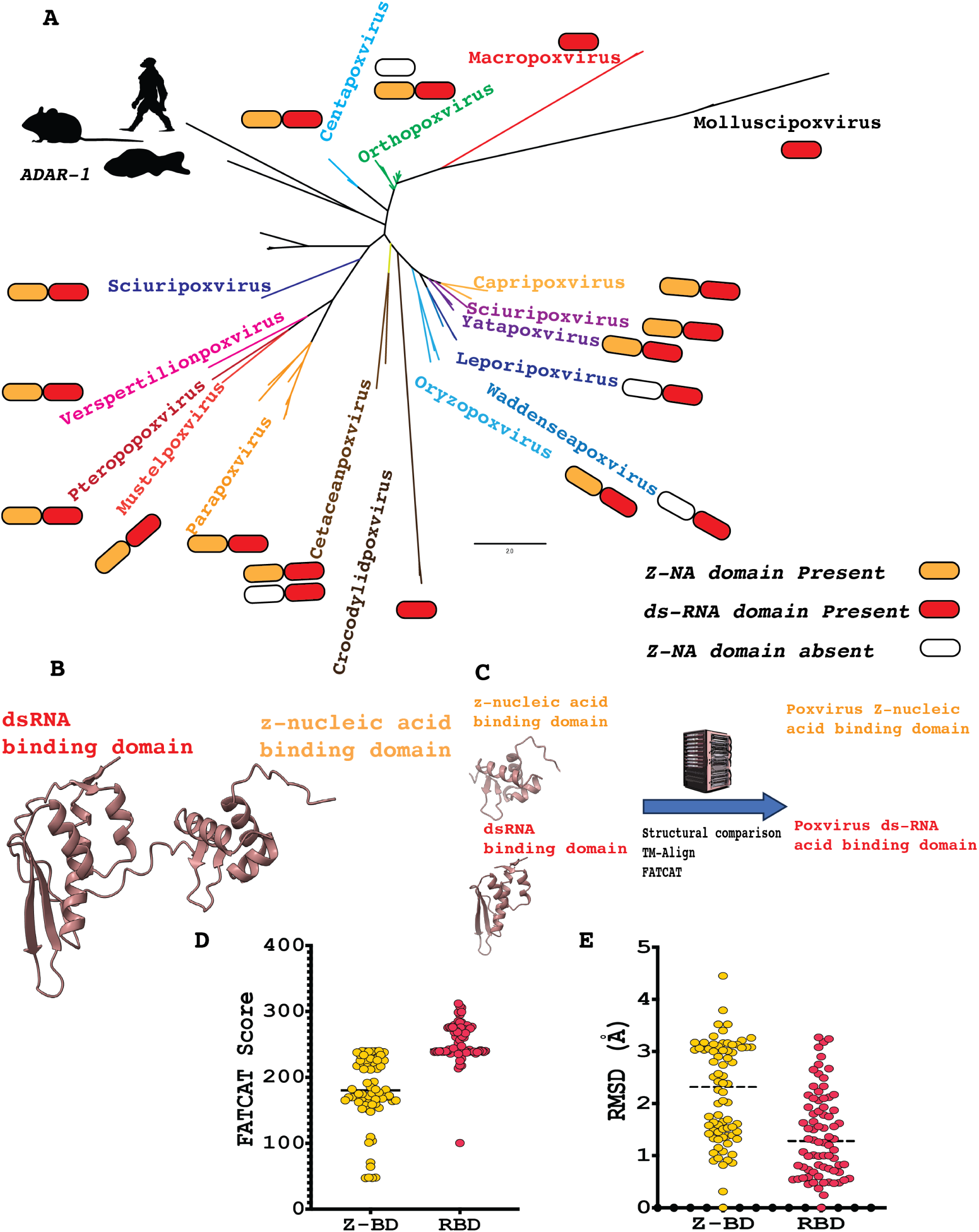
Recurrent loss of Zα-BD of E3-like proteins. (A) Phylogenetic tree of poxvirus dsRNA-BD containing proteins. Human, Mouse and Fish ADAR1 protein was used as the outgroup. (B) Alpha fold generated structure of VACV E3 protein depicting the dsRNA-BD (RBD) (red) and Zα-BD (yellow). (C) Figure depicting FATCAT and TM-Align structural comparisons of VACV Zα-BD and dsRNA-BD to poxvirus proteins containing homologues to these domains. (D) FATCAT score and (E) RMSD value of all poxvirus Zα-BD and dsRNA-BD compared to VACV E3 standard.

Leporipoxvirus family members (Myxoma virus and Shope Fibroma virus) and Waddenseapoxvirus lack an N-terminus Zα-NA domain, although structural comparison with VACV dsRNA-BD shows a significant structural similarity (Figures 3A-3C). The structural model of Myxoma virus (MYXV) E3-like protein M029 forms an alpha helix at the N-terminus that is not present in Shope Fibroma virus (SFV) gp029, its close cousin (Figures 3A and 3B). Wadden Sea poxvirus E3-like protein also lacks an N-terminus region (Figure 3C). Cetaceanpoxvirus contains two copies of the E3-like protein, one harboring a complete Zα-NA domain and the other missing the N-terminus region (Figure 3D).

**Figure 3.**
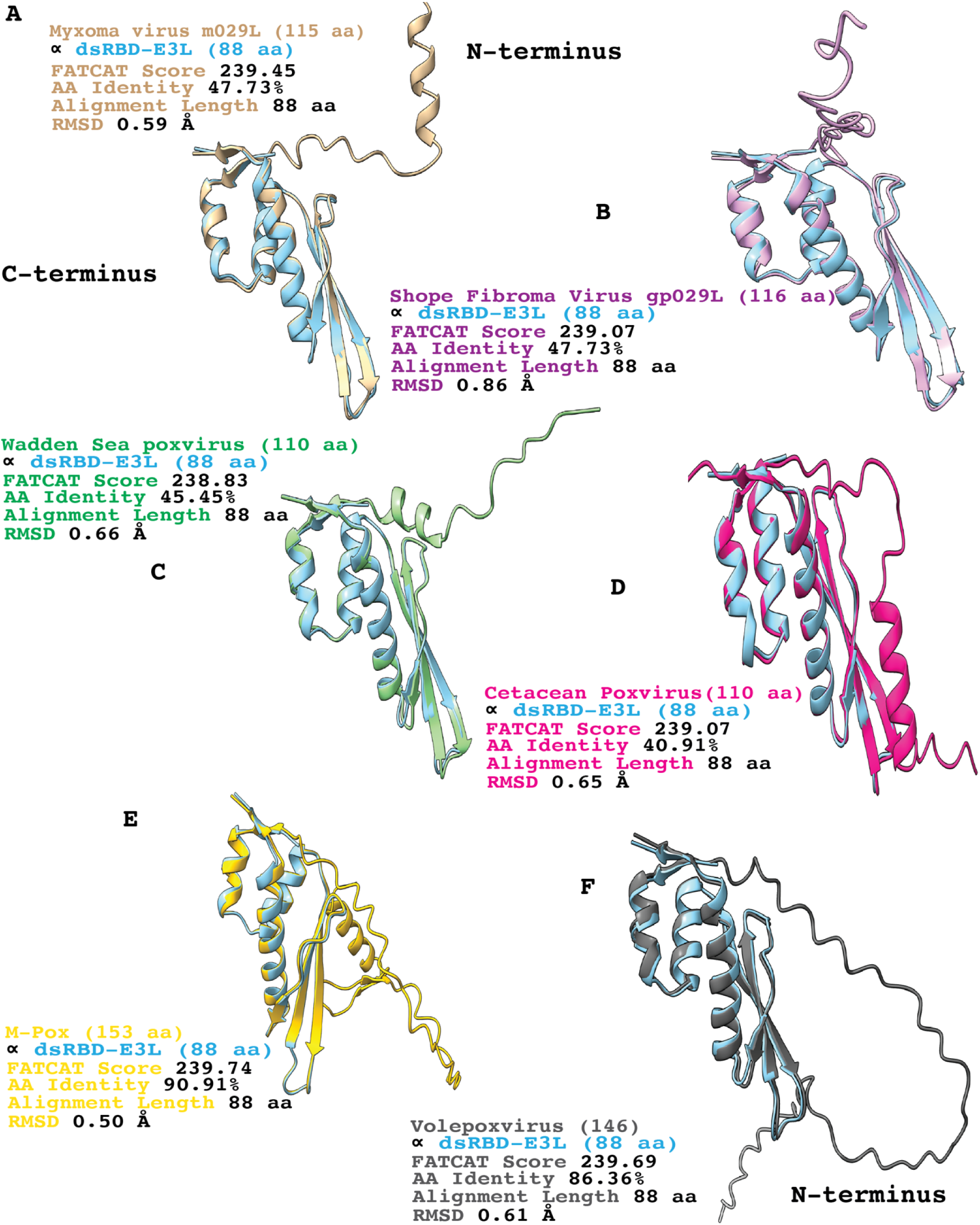
Recurrent loss of the Zα-BD amongst Chordopoxviruses. Structural homology showing FATCAT score, amino acid identity, alignment length and RMSD value comparing the dsRNA-BD of VACV to (A) Myxoma virus m029, (B) Shope Fibroma virus gp029, (C) Wadden Sea poxvirus E3-like proteins, (D) Cetacean poxvirus E3-like proteins, (E) M-Pox virus E3-like proteins and (F) Volepoxvirus E3-like proteins.

On the other hand, although most members of Orthopoxviruses possess an intact Zα-NA domain, some members are missing part (M-Pox) or the entire N-terminal region of the Zα-NA domain (Volepoxvirus) (Figures 3E and 3F).

### Leporipoxviruses activate necroptosis in necroptosis-competent human cells

Compared to Orthopoxviruses, such as VACV and CPXV, which have previously been shown to inhibit necroptosis and contain an E3 protein with an N-terminus Z-NA binding domain, it isn’t known whether different members of Leporipoxviruses induce necroptosis^4,8,19^. To investigate whether Leporipoxviruses activate necroptosis, we utilized two human cell lines, HT29 and Colo205, that are competent for necroptosis. We sequenced the transcriptome of HT29 cells in the presence and absence of human interferon beta to get a picture of the expression profile of the genes involved in necroptosis. Our data showed that MLKL and ZBP1, the host sensor of Z-nucleic acid, were IFN inducible (Figures S4A and S4B). Human HT29 cells were primed with human IFN-β for 24 hours and infected with representative members of Leporipoxviruses (MYXV-Lau, MYXV-Tol, and SFV) and Orthopoxviruses (VACV and CPXV) known to inhibit necroptosis. An hour before infection, cells were treated with a pan-caspase inhibitor z-VAD-FMK and infected with diverse Poxviruses. At 1-, 12-, 24-, and 36 hours post-infection, cells were treated with sytox orange and imaged using a Nikon Ti2 fluorescence microscope. At 24 hours post-infection, Leporipoxviruses (MYXV-Lau, MYXV-Tol, and SFV) had more sytox-positive cells than Orthopoxvirus (VACV and CPXV); however, VACV-Δ83N (VACV lacking the N-terminus Zα- BD), positive control for virus-induced necroptosis had the highest amount of sytox positive cells (Figure 4A). Viability measurements at 1-, 12-, 24-, and 36 hours after infection with MYXV-Lau, MYXV-Tol, and SFV showed less than 50% viable cells at 36hpi. Wild-type Cowpoxvirus had greater than 98% viable cells at 36hpi, values comparable to the untreated cells. However, wild-type VACV showed a marked 35 percent decreased viability compared to CPXV, while 99% of the cells died in the VACV-Δ83N infection condition at 36 hours (Figures 4A and 4B). We then monitored the level of MLKL phosphorylation to confirm necroptosis. HT29 cells were collected at 24hpi in the presence of phosphatase and protease inhibitors and subjected to western blot analysis. The results showed that MYXV-Lau, MYXV-Tol, SFV, and VACV-Δ83N infections led to the accumulation of phosphorylated MLKL (pMLKL), infection with WT-VACV resulted in a trace amount of pMLKL accumulation, whereas infection with CPXV produced little to no detectable MLKL phosphorylation (Fig. 4C). We repeated the same experiment in human Colo205 cells which are necroptosis competent and observed a similar pattern of cell viability and MLKL phosphorylation (Figure S4C and S4D). Thus, our data demonstrate that Leporipoxviruses activate necroptosis in human HT29 and Colo205 cells, suggesting that leporid-infecting poxviruses might not have a countermeasure to necroptosis.

**Figure 4.**
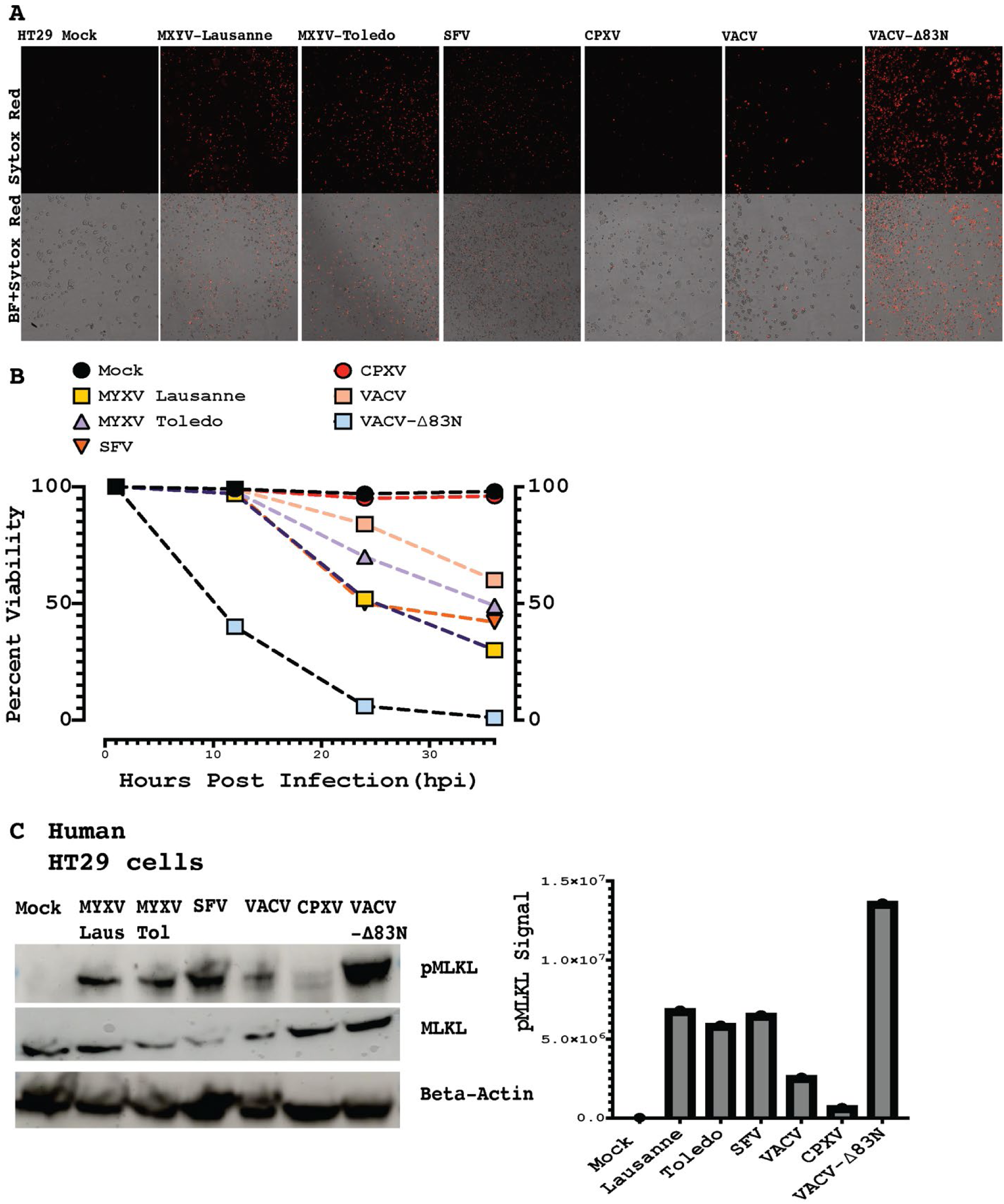
Leporipoxviruses stimulate necroptosis in necroptosis competent human cells. (A) Fluorescence images of interferon primed HT29 cells either uninfected or infected with MYXV-Lausanne, MYXV-Toledo, SFV, CPXV, VACV and VACV-Δ83N. Sample were stained with sytox orange dye and imaged 24 h.p.i. (B) HT29 cells viability of mock and virally infected samples collected at 1, 12, 24 and 36 hours after virus infection. (C) Western blot probing for phosphorylated MLKL, MLKL and beta actin levels in infected and uninfected samples and pMLKL signal intensity measured using imagestudio (Right panel).

### Myxoma virus activates both RIPK1 and RIPK3-mediated necroptosis in human cells

Since Leporipoxviruses have evolved in a species that are missing core executors of the necroptotic pathway, we wondered how key proteins of the necroptotic apparatus would behave in the presence of MYXV-Lau, a model leporipoxvirus. We used known inhibitors of these necroptosis proteins in human HT29 cells to test the effect of RIPK3, RIPK1, and MLKL on MYXV-mediated activation of necroptosis. We pretreated IFN-primed human HT29 cells with inhibitors of RIPK1 (GSK’963), RIPK3 (GSK’872), and Necrosulfonamide (NSA) in the presence of z-VAD FMK, or cells left untreated for one hour. Next, the cells were infected with MYXV-Lau, and sytox viability fluorescent images were taken at different time points. MYXV infection led to a greater than 60% decrease in cell viability at 36 hours post-infection (Figures 5A and 5B). However, inhibition of RIPK1 or RIPK3 increased cell viability compared to infection with virus alone at 36 hours.

**Figure 5.**
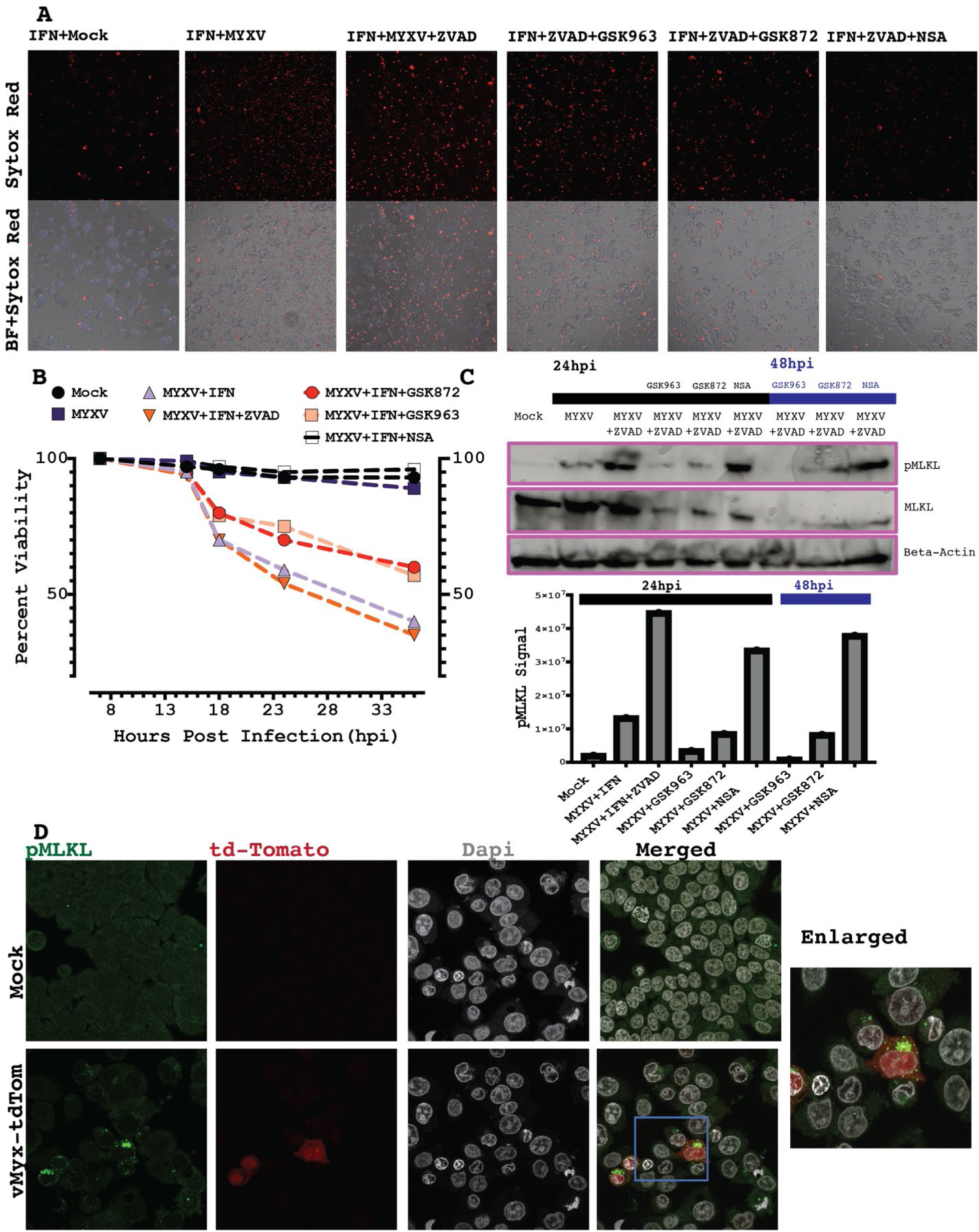
Myxoma virus activated necroptosis in necroptosis competent human cells. (A) Fluorescence images of interferon primed HT29 cells; mock uninfected, MYXV infected treated with ZVAD, MYXV virus infection in the presence of ZVAD and GSK963 or GSK872 or NSA. Samples were imaged 24 h.p.i in the presence of sytox orange. (B) Experiment A was repeated and viability measurement were collected at 7, 15, 18, 24 and 36 hours after MYXV infection. (C) Western blot of IFN-primed HT29 cells with the same treatments as in A probing for human pMLKL, MLKL and beta-actin, harvested at 24 and 48 h.p.i., with pMLKL intensity signals (bottom panel). (D) Confocal images of pMLKL in mock and myxoma virus infected IFN-primed HT29 cells, fixed at 24 hours after virus infection.

Surprisingly, inhibition of MLKL via Necrosulfonamide restored MYXV-infected cell viability to about 98%, 36 hours after infection (Figures 5A and 5B). Myxoma virus infection stimulated the accumulation of phosphorylated MLKL without z-VAD-FMK treatment, suggesting that caspase inhibition is not critical for MYXV-activated necroptosis. However, Z-VAD inhibition increased the amount of accumulated pMLKL (Figure 5C).

The inhibition of RIPK1 and RIPK3 reduced pMLKL accumulation, while treatment with NSA accumulated pMLKL even though it restored cell viability to greater than 98%. This is consistent with previous reports showing that MLKL is phosphorylated in the presence of NSA (Figure 5A- 5C)^20,21^. Immunofluorescence staining of pMLKL shows it localized to the cytoplasmic compartment and formed punctate structures 24 hours post MYXV infection (Figure 5D). Previous reports show that intracellular necroptosis depends on the accumulation of Zα-form nucleic acid^4^. We investigated whether MYXV infection accumulates Z-form nucleic acid in HT29 cells. HT29 cells were infected with either MYXV or Vacv-Δ83N for 24 hours and subjected to immunofluorescence staining. Samples were stained for both dsRNA and Zα-NA. Virus infection stimulated the formation of both Zα-NA and dsRNA, compared to uninfected cells (mock), although the distribution pattern between MYXV and VACV was different.

Interestingly, we observed a strong co-localization between the dsRNA and Zα-NA. The presence of dsRNA is mostly co-localized with Zα-NA; however, in some cells, Zα-NA was detected without dsRNA (Figure S5A). Taken together, our data demonstrate that in human necroptosis-competent HT29 cells, MYXV activates RIPK1 and RIPK3 necroptosis.

### RIPK1 and RIPK3 blockage rescues MYXV-activated necroptosis in necroptosis-competent mouse cells

Next, we tested whether MYXV infection can activate necroptosis in mouse cells and whether the components of necroptosis activation function like those of human cells. In this case, we used necroptosis-competent mouse L929 cells^4^. Mouse IFN-primed L929 cells were either left untreated or treated with z-VAD in the presence of GSK963, GSK872, and a PERK (Protein Kinase R (PKR)-like endoplasmic reticulum kinase) inhibitor, ISRIB. As previously observed in human cells, MYXV-activated necroptosis did not depend on caspase inhibition (Fig. 6A and 6C), suggesting that MXYV-encoded caspase inhibitors are functional in mouse-derived cells. At 12 hours post-infection, with or without interferon treatments, the viability of MYXV-infected L929 cells significantly reduced to only 25% (Figure 6B). RIPK1 inhibition via GSK’963 led to a significant increase in cell viability (more than 80% viable cells) compared to MYXV infection alone (Figure 6B). Additionally, inhibition of RIPK3 via GSK’872 increased cell viability (approximately 50% viable cells) compared to the virus-only conditions (Figure 6B). We confirmed that the cell death observed was due to necroptosis by performing a western blot analysis of phosphorylated MLKL. At 12 hours post-infection, western blot analysis shows that MYXV infection accumulates phosphorylated MLKL. However, z-VAD treatment reduced the strength of the pMLKL signal, while inhibition of RIPK1 and RIPK3 with GSK’963 and GSK’872 prevented the accumulation of pMLKL (Figure 6C). These results indicate that MYXV-mediated activation of necroptosis is functional in both human and mouse cells and depends on the critical components of the necroptosis machinery.

**Figure 6.**
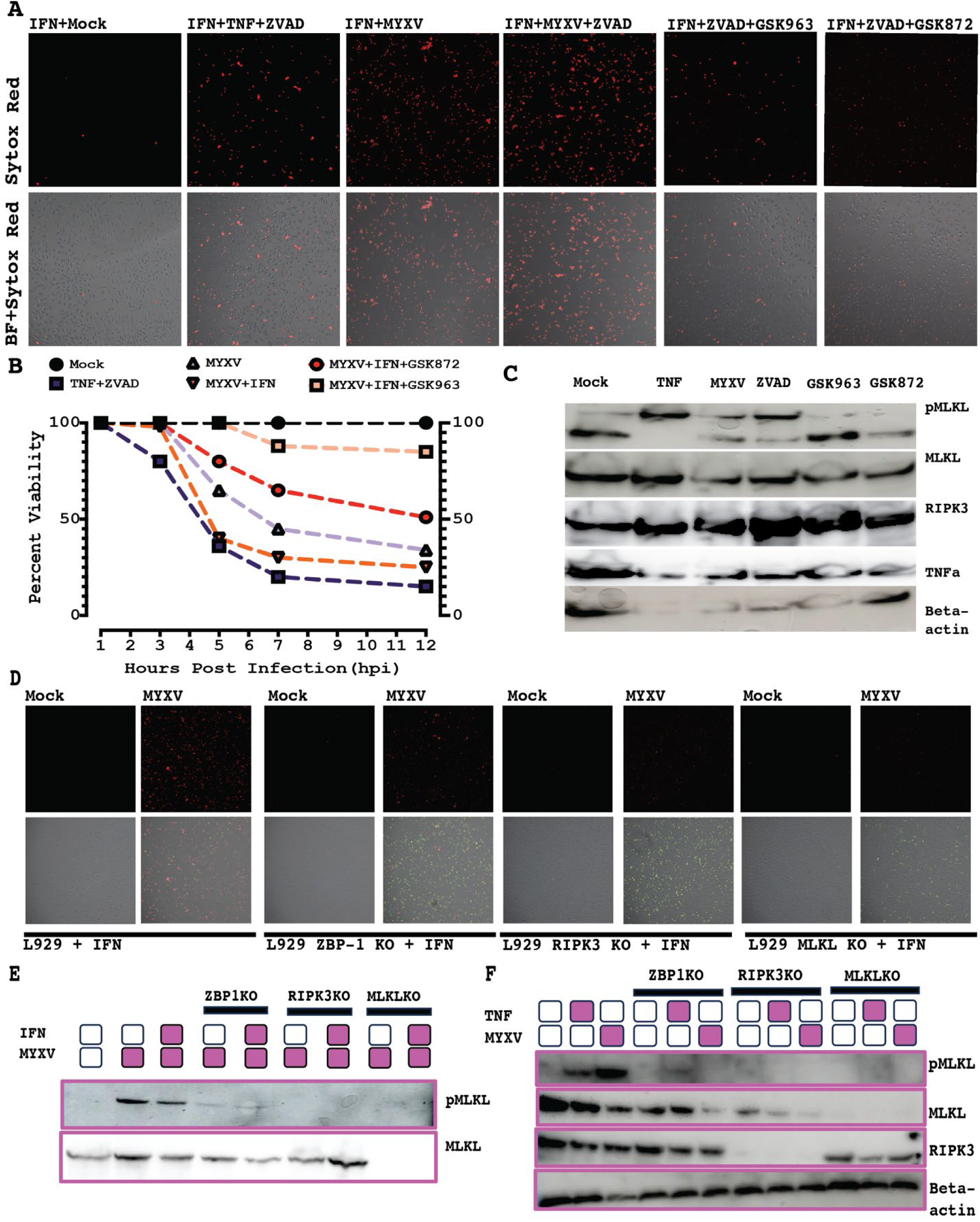
Inhibition of Necroptosis rescues MYXV replication in mouse L929 cells. (A) Fluorescence images of IFN-primed L929 cells with the following treatments: Untreated (Mock), ZVAD and TNF, MYXV, MYXV+ZVAD, MYXV+ZVAD+GSK963 and MYXV+ZVAD+GSK872. At 12 hours post treatment, cells were treated with sytox orange dye and images were captured. (B) Sample treatment was repeated as in A, and cell viability was calculated at 1, 3, 5, 7, and 12 hours after infection, by measuring cells that was positive of sytox related to the today cell number. (C) Western blot measuring the presence of pMLKL, MLKL, RIPK3, TNFa and beta-actin. Samples treatment were as decribed in A. (D) Fluorescence images of IFN-primed naïve L929, L929-ZBP1KO, L929-RIPK3KO and L929-MLKLKO cells, either left uninfected (mock) or infected with MYXV expressing GFP. (E) Western blot probing pMLKL in the presence and absence of IFN and MYXV, in the cell lines mentioned in figure D harvested at 24 hours after infection. (F) Western blot for pMLKL, MLKL, RIPK3 and beta-actin for IFN-primed samples treatments as in figure D, in presence of MYXV, TNF alpha, or no treatment.

### Activation of necroptosis inhibits MYXV replication in mouse L929 cells

To further understand the effect of various necroptotic components on MYXV replication, we used L929 cells that were genetically modified to lack ZBP1, RIPK3, or MLKL. To understand the effect of MYXV on cellular viability, we performed an MTS assay that measures cell viability via mitochondrial activity. At 24 hpi, the viability of unmodified naïve L929 cells dropped to about 25% (Figure S6A). The cell viability remained at 25% for the next several days that were tested (Figure S6A). ZBP1 knock-out cells showed a slower decrease in cell viability than naïve L929 cells. At 48 hpi, ZBP1KO L929 cells viability dropped to about 55% and remained at that level for several days. However, RIPK3 and MLKL knock-out L929 cells had minimal or no effect on cell viability after MYXV infection over the five-day course of the experiment (Figure S6A). Next, we quantified the impact of necroptosis-related genetic knock-outs on virus replication. ZBP1, RIPK3, and MLKL knockout in L929 cells increased myxoma virus titers–a 3-log increase on average. Compared to the L929 knock-out cell lines, naïve L929 cells showed little to no increase in viral titers (Figure S6B).

Since ZBP1, RIPK3, and MLKL deletion should affect necroptosis, we compared the effect of MYXV infection on naïve L929, along with L929 cells lacking either ZBP1, RIPK3, or MLKL. Cells were seeded in the presence or absence of IFN. IFN treatment progressed for 24 hours, and the cells were infected with MYXV expressing enhanced green fluorescence protein (gfp) and Tandem dimer Tomato protein (Tdtom). At 24 hours post-infection, the cells were harvested and subjected to western blotting. RIPK3 and MLKL knockouts had no MLKL phosphorylation on the blots (Figure 6E and S6C). We detected the phosphorylation of mouse MLKL in Naïve L929 cells in the presence and absence of IFN (Figure 6E). ZBP1 knockout cells had some trace amount of phosphorylated MLKL in the presence and absence of IFN, although not comparable to naïve L929 cells, suggesting that ZBP1 might not be the only pathway that feeds into MYXV-dependent necroptosis in mouse L929 cells (Figure 6E and S6C). We repeated the same experiment as before in the presence of IFN, and samples were collected at 8 hours post-infection and subjected to western blotting. We detected MLKL phosphorylation in naïve L929 cells with either MYXV or TNF treatment but not in the ZBP1, RIPK3, and MLKL knockout cells (Figure 6F). Sytox staining confirmed that ZBP1, RIPK3, and MLKL gene knock-out rescued MYXV virus-infected cell viability and replication (Figure 6D and S6B). Together, our data suggest that necroptosis inhibits myxoma virus replication in cells that are non-permissive to the virus. This indicates that MYXV does not contain a functional inhibitor of necroptosis.

### RIPK3 inhibition via GSK872 correlates with increased late viral gene expression in mouse cells

During this study, we observed that RIPK3 inhibition not only rescued myxoma virus-mediated cell death but also increased viral protein expression in mouse-derived cell lines at later time points of infection–24 and 48 hours post-infection. We treated L929 cells with GSK963, GSK872, and ISRIB in the presence of ZVAD and let the infection proceed for 48 hours. Cells were imaged for early myxoma virus gene expression (eGFP) and late viral gene expression (TdTom). Our fluorescence results show that RIPK3 inhibition had the highest fluorescence for the early and late genes (Figure 7A). We performed a Western blot analysis to detect TdTom expression and observed that RIPK3 inhibition had the highest TdTomato expression (late viral gene) (Figure 7B). We repeated the same experiment in mouse JC cells and observed a similar pattern of tdTomato expression in RIPK3- inhibited cells (Figure S7A). We did not observe a similar amount of myxoma virus gene expression in human-derived peripheral blood mononuclear cells (human PBMCs) upon RIPK3 inhibition.

**Figure 7.**
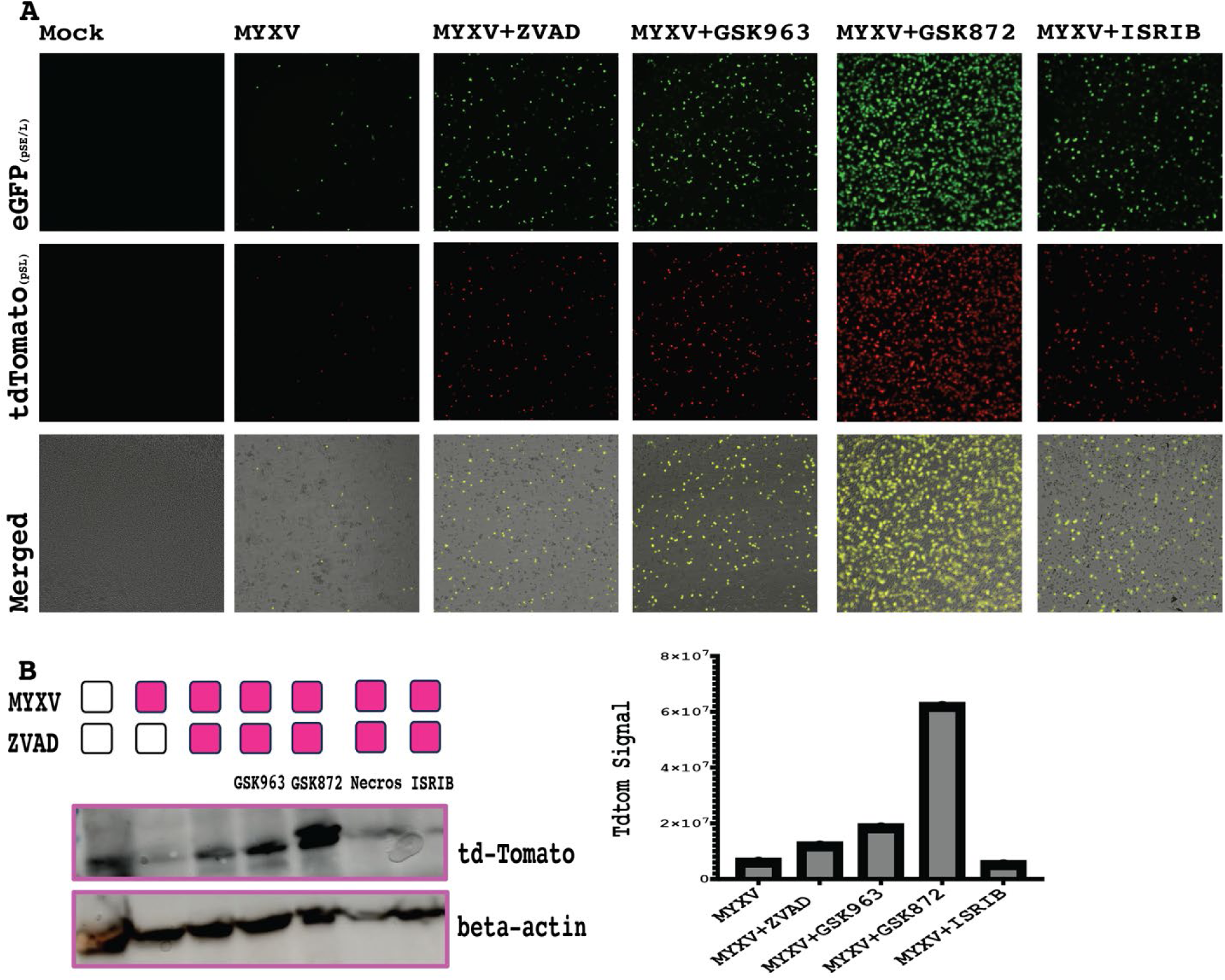
RIPK3 inhibition in mouse cells correlates with increased viral protein synthesis. (A) Fluorescence images L929 cells either uninfected (mock), or infected with MYXV, in the presence of ZVAD or/and GSK963, or GSK872 or ISRIB imaged 24 h.p.i. (B) Western blot measuring tdTomato expression in beta-actin in treatment conditions described in A. Signal intensity for tdTomato western blot (right panel).

This suggests that mouse RIPK3 affects virus gene expression and that mouse RIPK3 might affect viral protein synthesis.

## Materials and Methods

### Alpha Fold Singularity and Protein Structure Visualization

The E3-Like FASTA files for all poxviruses in our screen were obtained from (National Center for Biotechnology Information) NCBI. FASTA files were submitted to the AlphaFold2 singularity at the Arizona State University Research Computing Core. Computer-generated PDB files were visualized and overlayed using UCSF ChimeraX Daily, developed by the Resource for Biocomputing, Visualization, and Informatics at the University of California, San Francisco, with support from NIH P41-GM103311^22^.

Python scripts were implemented to run TM Align and FATCAT on our local supercomputing cluster^23,24^. RAW data was graphed on Graph Pad Prism.

### Phylogenetic Analysis

The E3L FASTA files were aligned using MUSCLE as implemented in MEGA 11^25^. All the E3L FASTA files in our screen were submitted to IQ-Tree 2 using the default settings to build a phylogenetic tree^26^. FigTree was used to visualize treefiles obtained from IQ-Tree^26^. FigTree files were visualized and illustrated using Adobe Illustrator.

### Synteny analysis

Genomes (genebank format) for representative poxviruses from each family were downloaded from NCBI. The files were trimmed to allow 10 genes on both sides of the dsRNA-BD-containing protein. Synteny analysis was performed using the default setting on clinker & clustermap.js^27^.

### RNA sequencing

Human HT29 cells were grown in 100 mm cell culture dishes. Samples were either treated with IFN or left untreated. Cells were harvested 24 hours post-treatment and subjected to RNA isolation.

RNA was sent to Novogene for RNA sequencing. Sequencing outputs were subjected to a standard data analysis pipeline that included differential gene expression and functional enrichment analysis using R. Gene count matrix file was used to create a heatmap of the genes involved in the establishment of necroptosis.

### Cells

Mouse L929, L929-ZBP1-KO, L929-RIPK3-KO, L929-MLKL-KO, JC, and JC-PKR-KO were obtained from the Jacobs lab at ASU. HT29 (catalog no. HTB-38), RK13 (catalog no. CCL-37), and Colo205 (catalog no. CCL-222) were purchased from ATCC (American Type Culture Collection). All the cell lines were tested for mycoplasma contamination before experimentation using the Universal Mycoplasma Detection Kit from ATCC (ATCC# 30-1012K). L929s were maintained in Minimum Essential Medium (MEM) (CORNING # 10-022-CV) supplemented with 5% FBS and/or 100ug penicillin-streptomycin (P/S; Cytiva# SV30010). HT29 was maintained in McCoy’s (Cytiva# SH30200.FS), 10% FBS, and 100ug P/S. Colo205 and JC cells were cultured in RPMI (Roswell Park Memorial Institute) (Cytiva# SG30027.02) with 10% FBS and 100ug P/S. Rabbit RK13 cells were maintained in Dulbecco’s Modified Eagle Medium (DMEM) (Cytiva # SG30022.LS) supplemented with 10% FBS and 100ug P/S. All cells were maintained at 37° Celsius in a 5% humidified incubator.

### Viruses

Myxoma virus Lausanne strain–vMyx-GFP-tdTomato (GFP is under the influence of the poxvirus synthetic early/late promoter (sE/L), and the gene for the tandem dimer tomato protein (tdTomato) is influenced by the poxvirus p11 late promoter)^28,29^. Myxoma virus Toledo (Toledo) has GFP expression and tdTomato under sE/L and p11 promoters, respectively^30^. We have previously published on Shope Fibroma virus expressing the lac-z gene^31^.

We used the Vaccinia virus Western Reserve strain, expressing sE/L GFP and late tdTomato^28^. Cowpox virus (CPXV) expressing sE/L GFP was reported previously by Lui et al^15^. These viral constructs were used, as described before^29^. Myxoma virus stocks were obtained through sucrose gradient purification^32^. Wild-type VACV and VACV with the N-terminus 83 amino acid deletion western reserve strain was obtained from Jacobs lab at Arizona State University and described previously^4^.

### Virus Titration

On the day of the experiment, cells were seeded in 24-well plates a day before infection and infected with the appropriate at moi. The virus was allowed to absorb for 1 hour and was later washed off with Phosphate-Buffered Saline (PBS) and replaced with fresh media. Cell pellet and supernatant were harvested at different time points after infection and frozen at -80 degrees. Samples were subjected to three freeze-thaw cycles and sonicated to release cell-associated virions. Samples were titrated in RK13 cells^32^.

### Antibodies

Human pMLKL ( Abcam # 187091, Cell Signaling # 91689S ), human MLKL (Abcam# ab243142, ab172868, ab184718, Cell Signaling # 14993S), human RIPK1 (Cell Signaling # 3493S), human phosphoRIPK1 (Cell Signaling # 65746S ), human RIPK3 (Cell Signaling # 13526), human phosphoRIPK3 (Cell Signaling # 57220S), human ZBP1 (Novus Biologicals # NBP1-7654), beta-actin(Life Technologies # MA5-15739-HRP), mouse pMLKL (abcam # ab196436, ), mouse MLKL (cell signaling # 37705S), mouse RIPK3 (Cell Signaling # 95702), mouse TNF-alpha (R&D Systems # AF-410-NA) and Z-NA ( Novus Biologicals # NB100-746)

### Western blotting and phospho-protein detection

For western blotting, protein samples were harvested, pelleted, and lysed using Radioimmunoprecipitation assay buffer (RIPA) lysis buffer in the presence of a protease-phosphatase inhibitors cocktail (ThermoFisher Scientific # 78446). Laemmli buffer (BIO-RAD #1610747) or SDS buffer containing 5% beta-mercaptoethanol was added to samples and heated at 95^0^ Celsius for 10 minutes and ran a Sodium Dodecyl Sulfate (SDS) protein gel. The SDS gel was transferred to a membrane and blocked with 1% Bovine Serum Albumin (BSA) in Tris-buffered saline with 0.1% Tween (TBST) buffer for one hour. Next, the blot was incubated overnight with the primary antibody, washed with TBST for 5 minutes 3 times, and incubated with a secondary antibody conjugated to HRP. The membrane was washed with TBST to get rid of the secondary antibody. Membranes were visualized using an Amersham ImageQuant^TM^ (Cytiva) 800 Western blot imager.

### Cell treatment and Sytox cell viability assay

Cells were seeded the day before the experiment, and treated with media containing and/or 100 ug mL^-1^ Benzyloxycarbonyl-Val-Ala-Asp(OMe)-fluoromethylketone (Z-VAD-FMK) (ApexBio # 1902), 2 ug mL^-1^ 1-[(5S)-4,5-Dihydro-5-phenyl-1H-pyrazol-1-yl]-2 (GSK’963) (Sigma-Aldrich # SLM2376- 25MG), 2 ug mL^-1^ N-(6-(isopropylsulfonyl)quinolin-4-yl)benzo[d]thiazol-5-amine hydrochloride (GSK’872) (Sigma-Aldrich # 5303890001) and 2 ug mL^-1^ Necrosulfanamide (EMD Millipore #480073) and 1ug mL^-1^ of sytox orange (Invitrogen #S11368). Samples were observed using the EVOS live cell imaging system, and representative images from different time points were quantified by counting the dead cells that contained the sytox dye compared to the live cells that incorporated the nuclei stain Hoechst (Thermo Scientific^TM^ #62249).

### MTS cell viability assay

Cells were seeded in a 96-well plate to a 90% confluency and infected with vMyx-GFP-tdTomato . At the appropriate time points, the MTS solution (3-(4,5-dimethylthiazol-2-yl)-5-(3- carboxymethoxyphenyl)-2-(4-sulfophenyl)-2H-tetrazolium) (Promega # G3581) was added to individual wells. Samples containing MTS solution were incubated for 1 hour, and the absorbance at 490 nm was recorded using a microwell plate reader (Thermo Scientific^TM^ Varioskan LUX multimode microplate reader). Time points were recorded in triplicates.

### Immunofluorescence assay and fluorescence imaging

Cells were seeded overnight on an 8-well Ibidi dish and infected with different viruses. At the appropriate time points, samples were fixed using paraformaldehyde (PFA) for 5 minutes, rinsed with PBS, and permeabilized using triton-X 100 for 2 minutes. After permeabilization, the sample was fixed with PFA for 5 minutes, and blocked in 1% BSA for 1 hour at room temperature. The samples were incubated with the primary antibody overnight and washed with PBS supplemented with 0.1% tween (PBST). The secondary antibody conjugated to the fluorophore (Thermo Scientific^TM^ Alex Fluor 488, 568, and 647) was incubated for 1 hour at room temperature. The secondary antibody was washed off using PBST, and samples were immersed in PBS-containing 1ug/mL of 4′,6-diamidino-2-phenylindole (DAPI) and imaged using a Nikon AX confocal microscope at the Biodesign Imaging Core Facility.

Samples with varying treatments were imaged using a Nikon Eclipse Ti2 microscope (Nikon). The images were converted using Nikon software.

### Data Analysis and Figure Design

All experiments were performed at least three times independently. A 2-way ANOVA and multiple comparisons in Graph Pad Prism software were used to compare different treatments for statistical comparisons. All figures were designed using Adobe Illustrator, except the graphical abstract, created using Biorender.

## Discussion

In this study, we defined the origin of E3-like proteins within Poxviruses. Our data supports that poxviruses contain 3 unique categories of dsRNA-BD containing proteins acquired independently during evolution. Structural comparison using alpha-fold of the core dsRNA-BD to that of vaccinia virus E3 showed significant similarity within all 3 categories of dsRNA-BD-containing proteins, suggesting that dsRNA-BD-containing proteins are core immune modulators of poxvirus infection. Traditionally, based on protein sequence homology, it was known that molluscipoxviruses lacked an E3 orthologous protein that regulates PKR, possibly due to high sequence divergence between MC141 protein and E3-like proteins^7,33^. Our structural homology comparison for the first time shows that macropoxvirus and molluscipoxvirus contain a dsRNA-BD-containing protein.

Nonetheless, using the same approach, we could not find a dsRNA-BD containing protein for two Chordopoxviruses (Salmonpoxvirus and Avipoxviruses) (Figure 1A).

We previously reported that certain mammalian lineages are missing key proteins in the necroptosis pathway, and their accompanying poxviruses lack the N-terminus Z nucleic acid binding domain, critical for necroptosis inhibition. In this work, we identified members from four different families of poxviruses that are fully or partially missing the N-terminus, Zα-BD: Leporixpoxvirus (Myxoma virus, Myxoma virus Toledo, Shope Fibroma virus), Waddenseapoxvirus, Cetaceanpoxvirus and Orthopoxvirus (M-Pox and Volepox) (Figure 2A). Recurrent Zα-BD loss amongst different poxvirus families seems to be an independent event (Figure 2A, S1A). Interestingly, we observed that the N-terminus domain of the E3-like homologous proteins seems to be an additional secondary structure when compared to the canonical Zα-BD in the Vaccinia virus (Figure 2D).

However, this was not the case for the dsRNA-BD (Figure 2D). These observations suggest that the dsRNA-BD might be under higher selective pressure than the Zα-BD. However, we do not yet have a good theoretical explanation for the recurrent loss and extensive modifications at the N-terminus domain of E3.

The structures of the E3-like proteins missing their Zα domain have a FATCAT score greater than 238 when compared with the dsRNA-BD of Vaccinia virus (Figure 3A-F). This suggests conservation in the dsRNA-BD domain. On the other hand, the N-terminus modification seems to be unique for each poxvirus, suggesting a varying degree of adaptation at the N-terminus domain. In all, poxviruses lacking an N-terminus Zα-BD contain unique secondary structures at the N-terminus that might have adapted for diverse host-specific functions from Zα-Nucleic acid regulation.

In this work, we compared selected members of Orthopoxviruses, known to inhibit necroptosis, with Leporipoxvirus, which we hypothesize would stimulate the necroptosis pathway because the host species in which they have evolved in lack key regulators of necroptosis^4,19,34,35^. In other words, Leporipoxviruses contain no functional inhibitors of necroptosis. Our data suggest that in human necroptosis-competent cells, Leporipoxviruses promoted cell death and the accumulation of pMLKL to levels greater than wild-type Orthopoxviruses. This suggests that Leporipoxvirus are naked to necroptosis or possibly do not encode any functional inhibitors of the pathway (Figure 5). Although Leporipoxvirus activated necroptosis to levels greater than wild-type Orthopoxviruses, Vaccinia virus missing the Zα-promoted rapid cell death and more significant amount of pMLKL than any of the Leporipoxviruses. Thus, Leporipoxviruses did not phenocopy VACV-Δ 83 N in terms of activation of necroptosis, even in the absence of Zα-NA-BD. We think the higher cell death observed with VACV-Δ 83 N is because Orthopoxviruses replicate at higher kinetics than Leporipoxvirus in human and mouse-derived cells.

In both human and mouse-derived necroptosis-competent cells, we observed that RIPK1, RIPK3, and MLKL all contribute to the myxoma virus-induced necroptosis to varying degrees. Treatment with both RIPK1 and RIPK3 inhibitors partially rescued MYXV-induced necroptotic cell death.

However, MLKL inhibitor NSA almost entirely rescued cell death. These observations support that RIPK1 and RIPK3 are involved in the MYXV-induced activation of necroptosis. Koehler et al. previously demonstrated that the Vaccinia virus lacking the Zα-NA binding domain activates RIPK3, not RIPK1 necroptosis^4^. These differences between MYXV and VACV suggest that there are possibly other inhibitors for the necroptosis pathway present in VACV but lacking in MYXV. Apart from poxviruses, many viruses and their proteins target RIPK1, RIPK3, and MLKL to inhibit necroptosis^36^. For example, herpesvirus-encoded RHIM proteins M45, ICP6, and ICP10 impede the formation of RIPK3 amyloid; and Epstein-Barr virus (EBV)-encoded latent membrane protein 1 (LMP1) blocks the formation of RIPK1 and RIPK3 necrosome. Using MYXV, we demonstrate that viruses without any functional inhibitors for necroptosis can be used as a model to understand the role of different components of the necroptosis pathway.

Our studies demonstrate that the functional necroptosis pathway inhibits MYXV replication and the formation of infectious progeny virions. This is possibly one of the mechanisms of host restrictions of MYXV. Therefore, it is tempting to speculate if MYXV can cause myxomatosis in rabbits with functional necroptosis machinery.

Necroptosis is an inflammatory form of cell death that has been shown to have varying effects on cancer progression, metastasis, and clearance^34,37–39^. Myxoma virus is currently being developed as an oncolytic virus for cancer treatment^40,41^. It would be interesting to compare the effect of necroptosis on cancer therapy, where a virus that naturally triggers necroptosis versus a virus that has evolved to shut down necroptosis and most immune pathways.

---

**Figure S1.**
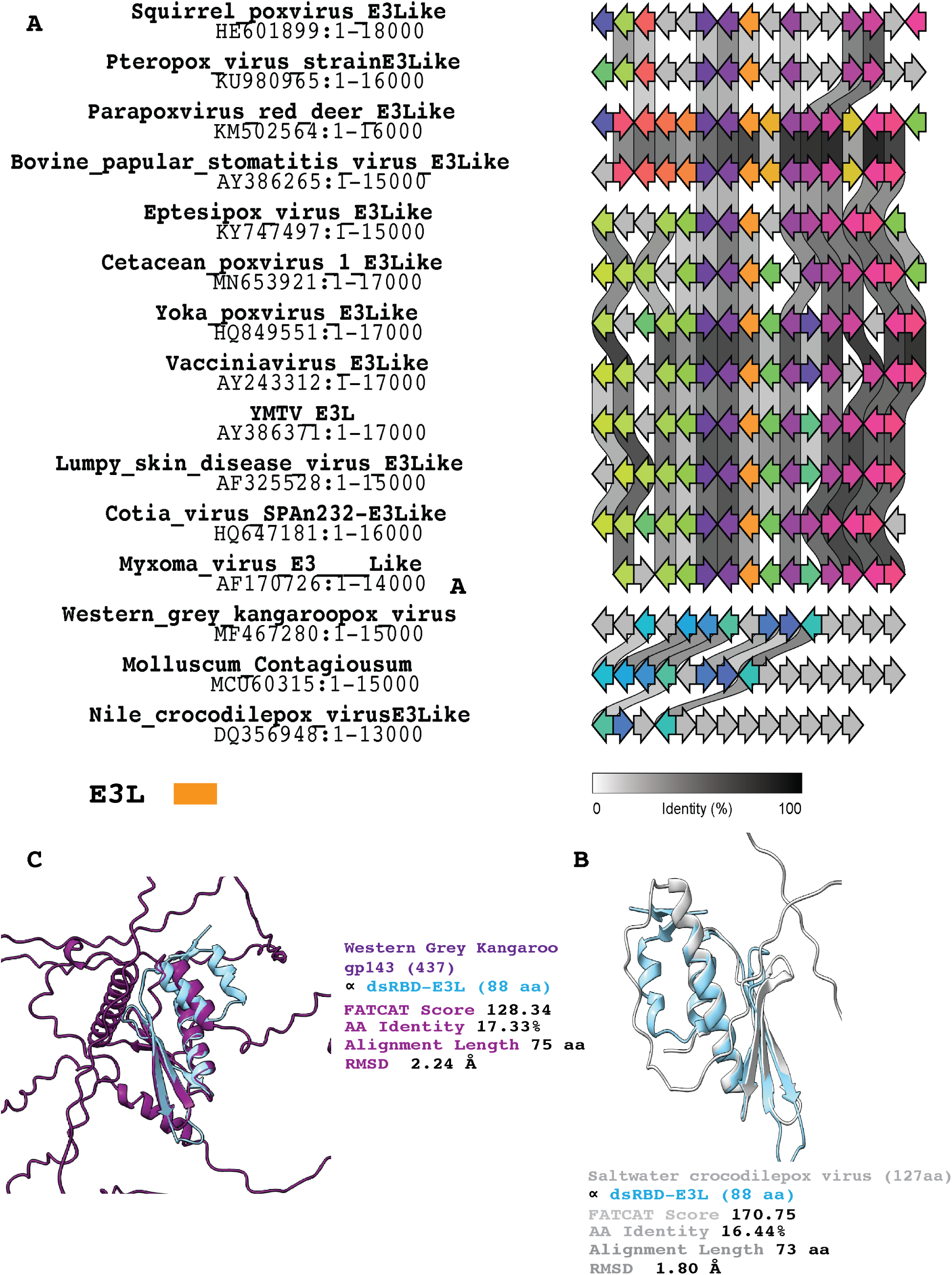
Origin of Poxvirus dsRNA-BD containing proteins. (A) Synteny analysis showing the relatedness of the genes surrounding Poxvirus dsRNA-BD containing proteins. Protein structural homology showing FATCAT score, amino acid identity, alignment length and RMSD value comparing the dsRNA-BD of VACV to (B) Saltwater Crocodilepoxvirus and (C) Western Grey Kangaroo poxvirus gp143.

**Figure S4.**
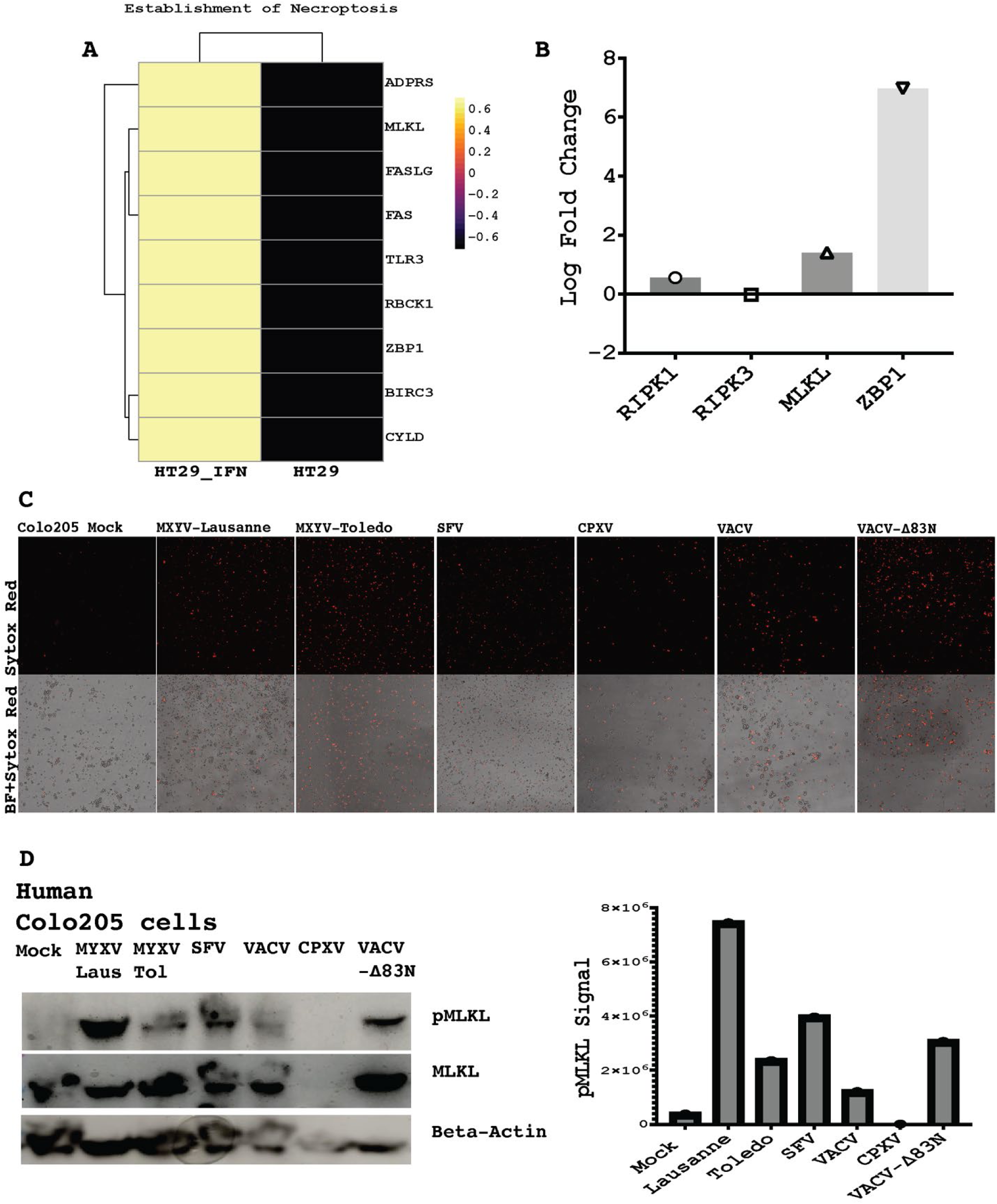
Leporipoxviruses activate necroptosis in necroptosis competent human cells. (A) Heatmap depicting differentially expressed genes involved in establishing necroptosis and (B) Log fold change in RIPK1, RIPK3, MLKL and ZBP-1 in human HT29 cells. (C) Fluorescence images of interferon primed Colo205 cells either uninfected or infected with MYXV-Lausanne, MYXV-Toledo, SFV, CPXV, VACV and VACV-Δ83N. Sample were stained with sytox orange dye and imaged 24 h.p.i. (D) Western blot probing for pMLKL, MLKL and beta actin levels in infected and uninfected IFN-primed Colo205 cells. pMLKL signal intensity was measured using imagestudio (Right panel).

**Figure S5.**
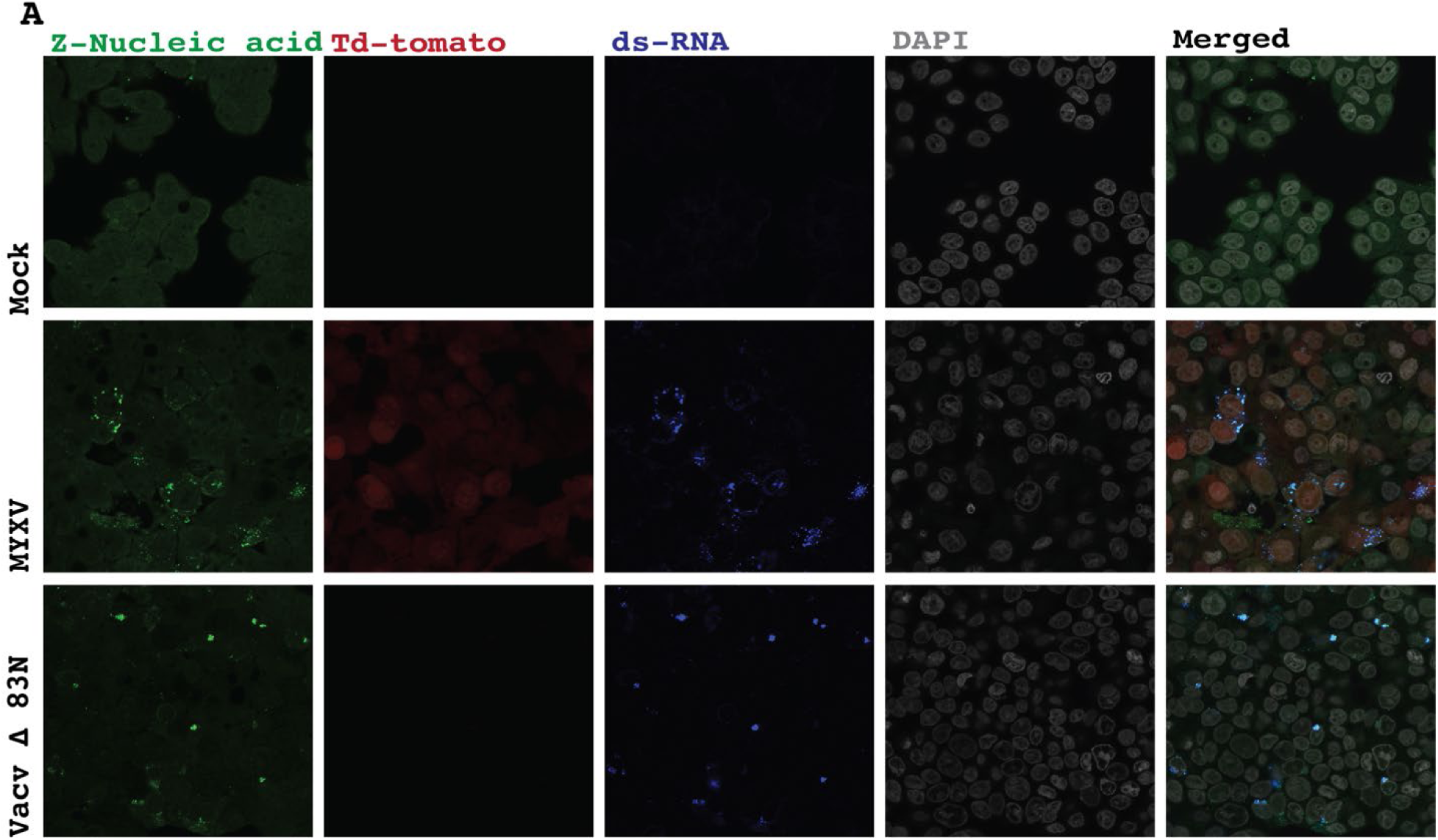
Myxoma virus infection accumulates Zα-form nucleic acid. (A) Confocal images of Z-Nucleic acid and ds-RNA accumulation of unfected, MYXV and VACV-Δ83N infected IFN-primed HT29 cells, fixed at 24 hour after virus infection.

**Figure S6.**
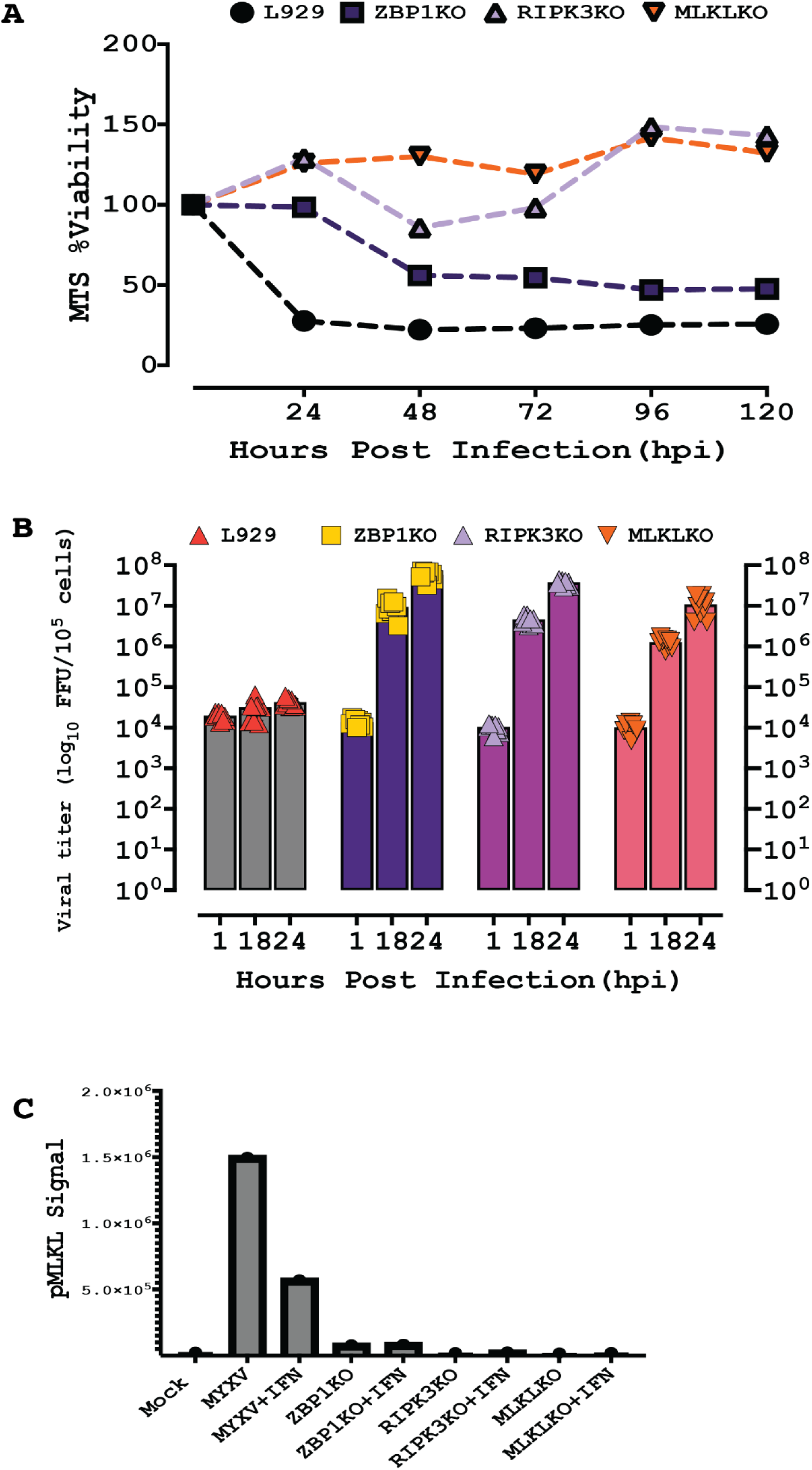
Necroptosis inhibits MYXV replication in mouse L929 cells. (A) MTS assay measuring mitochondria activity at 1, 24, 48 96 and 120 hours after virus infection in naïve L929, L929-ZBP1KO, L929-RIPK3KO and L929-MLKLKO cells. (B) MYXV titration in naïve L929, L929-ZBP1KO, L929-RIPK3KO and L929-MLKLKO cells at 1, 18 and 24 h.p.i. (C) pMLKL signal intensity for MYXV infected naïve L929, L929-ZBP1KO, L929-RIPK3KO and L929-MLKLKO cells in the presence and absence of IFN.

**Figure S7.**
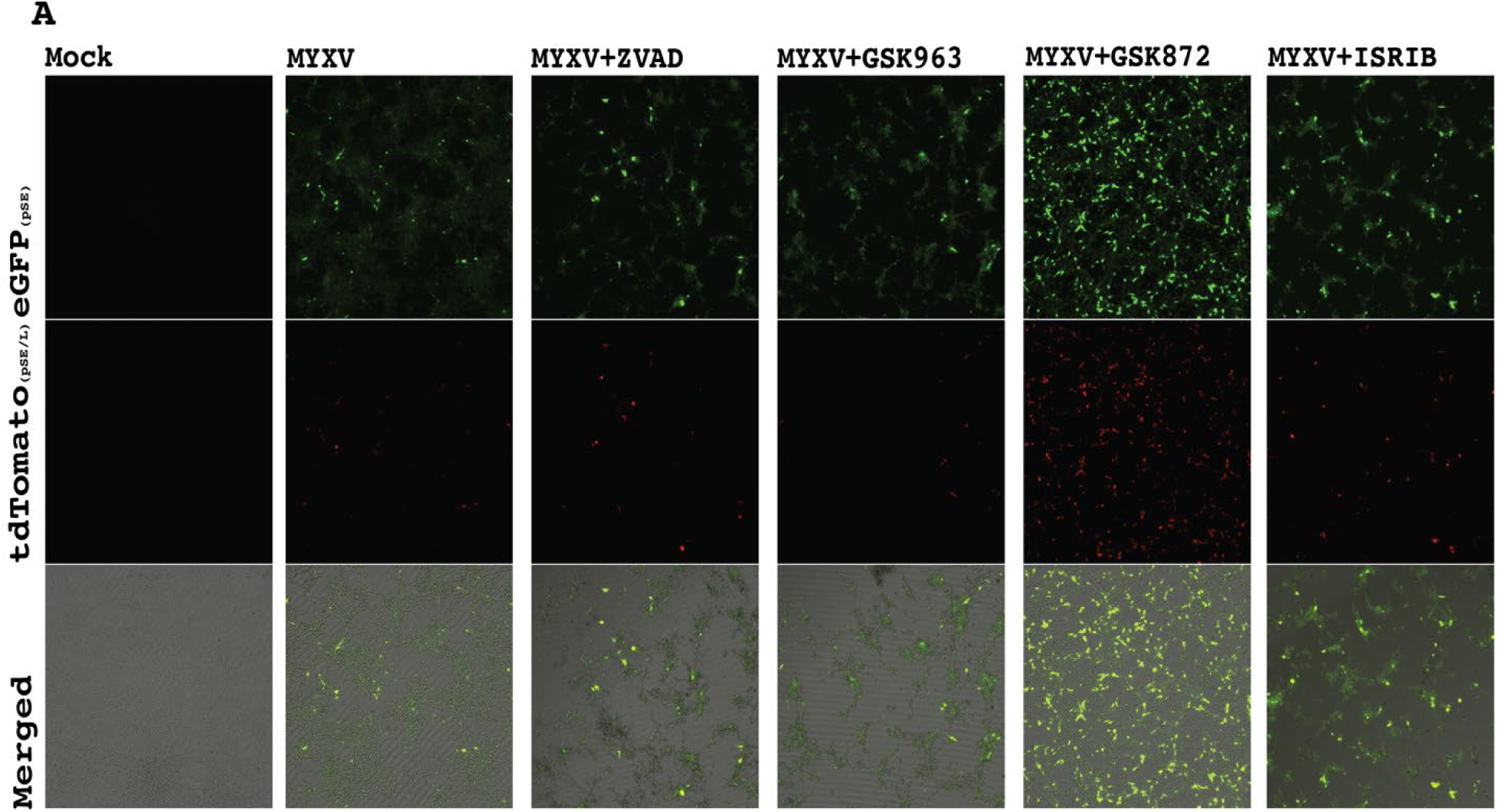
RIPK3 inhibition upregulates MYXV early and late gene expression. (A) Fluorescence images of mouse JC cells with the following treatments: mock (uninfected), MYXV, MYXV and ZVAD, MYXV in the presence of ZVAD and GSK963 or GSK872 or ISRIB.

